# A single base pair substitution on Chromosome 25 in zebrafish distinguishes between development and acute regulation of behavioral thresholds

**DOI:** 10.1101/2023.08.25.554673

**Authors:** Elelbin A. Ortiz, Philip D. Campbell, Jessica C. Nelson, Michael Granato

**Affiliations:** Department of Neuroscience, University of Pennsylvania; Department of Cell and Developmental Biology, University of Pennsylvania; Department of Psychiatry, University of Pennsylvania; Department of Cell and Developmental Biology, University of Colorado Anschutz Medical Campus.

## Abstract

Behavioral thresholds define the lowest stimulus intensities sufficient to elicit a behavioral response. Establishment of baseline behavioral thresholds during development is critical for proper responses throughout the animal’s life. Despite the relevance of such innate thresholds, the molecular mechanisms critical to establishing behavioral thresholds during development are not well understood. The acoustic startle response is a conserved behavior whose threshold is established during development yet is subsequently acutely regulated. We have previously identified a zebrafish mutant line (*escapist*) that displays a decreased baseline or innate acoustic startle threshold. Here, we identify a single base pair substitution on Chromosome 25 located within the coding sequence of the *synaptotagmin 7a* (*syt7a*) gene that is tightly linked to the *escapist* acoustic hypersensitivity phenotype. By generating animals in which we deleted the *syt7a* open reading frame, and subsequent complementation testing with the *escapist* line, we demonstrate that loss of *syt7a* function is not the cause of the *escapist* behavioral phenotype. Nonetheless, *escapist* mutants provide a powerful tool to decipher the overlap between acute and developmental regulation of behavioral thresholds. Extensive behavioral analyses reveal that in *escapist* mutants the establishment of the innate acoustic startle threshold is impaired, while regulation of its acute threshold remains intact. Moreover, our behavioral analyses reveal a deficit in baseline responses to visual stimuli, but not in the acute regulation of responses to visual stimuli. Together, this work eliminates loss of *syt7a* as causative for the *escapist* phenotype and suggests that mechanisms that regulate the establishment of behavioral thresholds in *escapist* larvae can operate largely independently from those regulating acute threshold regulation.

## Introduction

Behavioral thresholds are critical for organisms to respond to relevant environmental cues or threats, while ignoring irrelevant environmental stimuli(1–3). A prime example of this is the acoustic startle response, a defensive and highly conserved behavior across vertebrates that exhibits a quantifiable behavioral threshold(4). Across species, organisms establish innate baseline threshold which are shaped during development(5–7). Baseline startle thresholds can subsequently be modulated acutely through sensory stimulation (e.g. through habituation or sensitization), yet provided with enough time will return to the baseline threshold. Differences in baseline startle thresholds across genetic strains as well as evidence of generational inheritance strongly suggest the existence of genetic components that establish baseline thresholds(8,9). Our current knowledge regarding the molecular and circuit mechanisms underlying the regulation of the startle threshold largely stems from a wealth of studies focusing on acute regulation, such as during short term habituation and pre-pulse inhibition(10–13). Dysregulation of both acute modifications and developmental mechanisms have been associated with several neuropsychiatric and neurodevelopmental disorders, such as in schizophrenia, autism spectrum disorders and anxiety disorders(14–17). While there has been significant progress in understanding the molecular mechanisms that acutely regulate the startle response(9,18), the molecular and circuit mechanisms underlying the establishment of the innate threshold of the acoustic startle response are only partially understood.

Zebrafish are a powerful system to identify molecular and circuit mechanisms that regulate behavioral thresholds. By 5 days post fertilization (5dpf), zebrafish larvae exhibit a wide variety of well characterized behaviors, including the acoustic startle response(7,19). Additionally, forward genetic screens have identified genes that regulate behavioral thresholds to acoustic and visual stimuli, emphasizing the power of this system to dissect genetic and molecular mechanisms of disease-relevant behaviors(10,18,20,21). Finally, cell types that mediate the initiation of the startle response have been identified, providing a coherent framework to explore mechanisms that distinguish between innate and acutely regulated thresholds(22,23).

Here we examine the process of establishing behavioral thresholds using *escapist*, a zebrafish mutant line identified in a forward genetic screen based on its hypersensitivity to acoustic stimuli(18). Using RNA sequencing we uncovered a single candidate gene, *synaptotagmin 7a*, linked to the *escapist* acoustic hypersensitive phenotype. Although tightly linked to the mutant phenotype, thorough behavioral analysis in combination with molecular genetic testing strongly suggests that loss of *synaptotagmin 7a* does not cause the *escapist* acoustic hypersensitive phenotype. However, through our extensive behavioral analysis, we uncovered an additional behavioral phenotype in *escapist* mutants. Specifically, we find that mutants exhibit reduced motor responsiveness to dark flashes, a stereotyped behavioral response to sudden darkness. Finally, we find that unlike the innate threshold, acute threshold modulation of visual and acoustic stimuli in *escapist* mutants is unaffected, consistent with the idea that mechanisms that regulate the establishment of innate thresholds can operate largely independently from those regulating acute thresholds.

## Results

### Identification of candidate genes underlying the *escapist* acoustic startle hypersensitivity phenotype

By 5dpf, zebrafish larvae respond to sudden and intense acoustic stimuli with a stereotypic short latency turn known as the startle response(24). The probability of the all-or-none startle response increases as the acoustic stimulus intensity increases(18). We previously performed a forward genetic screen and identified five zebrafish mutant lines that exhibit hypersensitivity of the acoustic startle response(18). To compare the strength of the hypersensitivity phenotype across wildtype and mutant animals we exposed larvae to various stimulus intensities and plotted the larvae’s probability to perform a short latency startle response (SLC) based on stimulus intensity. We define the area under the resultant curve as the sensitivity index(18). Specifically, for the *escapist* mutant line, we established crosses between *escapist* heterozygous carriers and divided the offspring larvae into two groups based on their sensitivity indices: putative mutants (top 25%) and putative siblings (bottom 75%). When compared to the top 25% most responsive larvae obtained from wild type incrosses, putative *escapist* larvae have significantly increased sensitivity (Figure 1A, 1B). Importantly, the bottom 75% (least responsive) escapist larvae are within range of the responsiveness of all wildtype larvae, consistent with recessive inheritance of the hypersensitivity phenotype (Figure 1A,1C).

**Figure 1.**
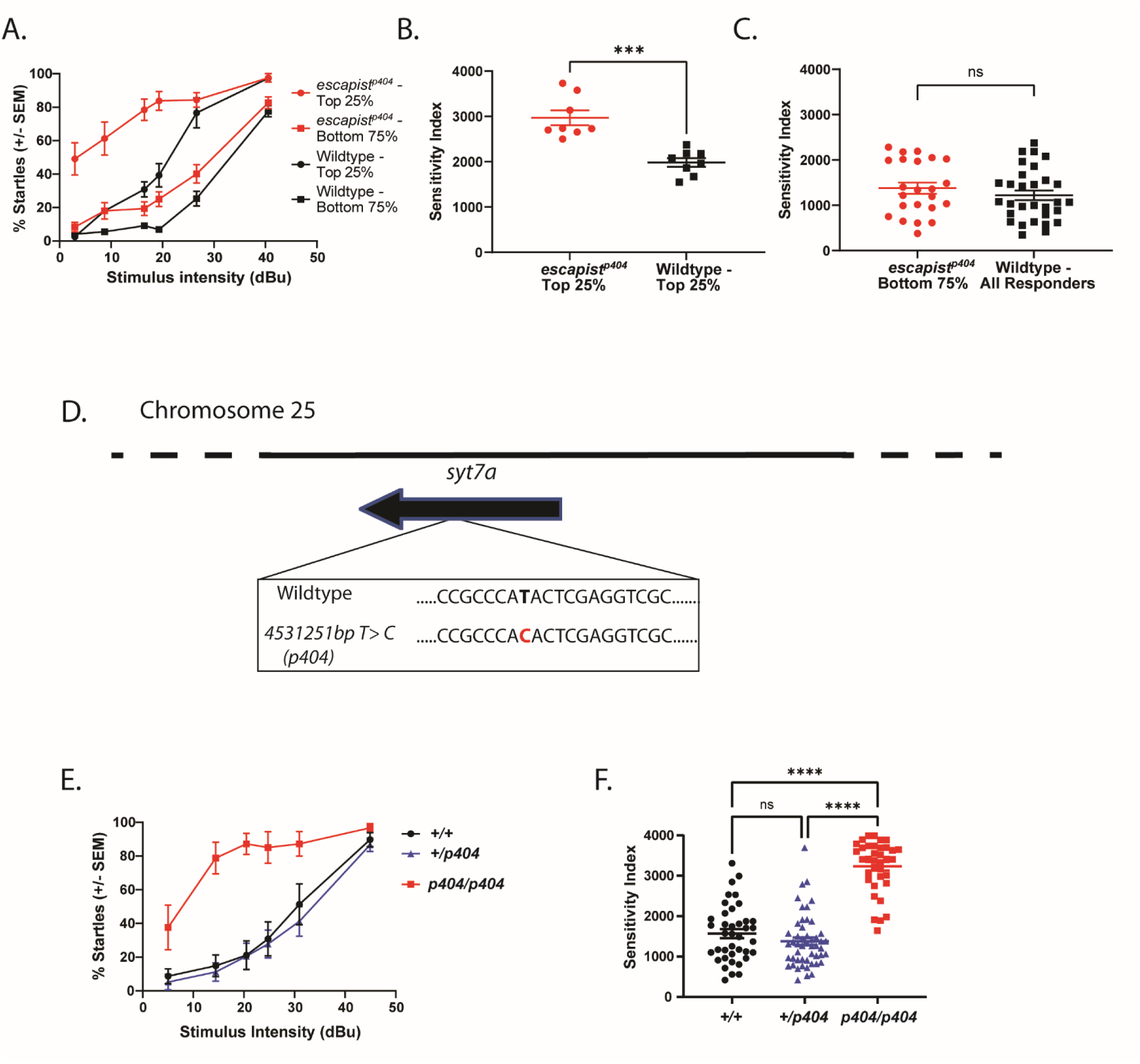
RNA sequencing linkage analysis identifies a single base pair change on Chromosome 25 that is tightly linked to the *escapist* hypersensitive phenotype. A) Acoustic sensitivity curve for *escapist* and wildtype (WIK) 5-day old larval zebrafish. X-axis represents stimulus intensity (dBu) and Y-axis represents percent short latency (SLC) startles. Larvae were placed in the following groups: the top 25% responsive larvae from the *escapist* line (*escapist-* Top 25%, n=8 larvae), the bottom 75% responsive larvae from the *escapist* line (*escapist* -Bottom 75%, n=23 larvae), the top 25% responsive larvae from the wildtype WIK line (Wildtype-Top 25%, n=8 larvae), the bottom 75% responsive larvae from the wildtype WIK line (Wildtype-Bottom 75%, n= 22 larvae) B) The area under the sensitivity curve was computed to generate the sensitivity index for each individual larva represented in Figure 1A. Sensitivity index for the top 25% responders from the *escapist* line (red circles) and wildtype WIK lines (black squares). Unpaired t-test for *escapist* Top 25% vs wildtype Top 25%: p=0.0001 C) Sensitivity index for the bottom 75% *escapist* responders (red circles) and all responders from the wildtype WIK lines (black squares). Unpaired t-test for *escapist* Bottom 75% vs wildtype All responders: p=0.3380 (ns) D) Schematic of linked region on Chromosome 25 identified through RNA Sequencing linkage analysis using zv9 as a reference genome. Analysis of linked mutations identified one single nucleotide base pair change (*chr25:4531251bp T->C* or *p404*) as highly linked to the *escapist* hypersensitive phenotype. Sequence shown is of the reverse strand. E) Acoustic sensitivity curve for 5dpf larvae from crosses of *escapist* carriers genotyped for the *p404* lesion. Homozygous wildtype (+/+, n=39 larvae) curve shown in black, heterozygous (+/*p404*, n= 50 larvae) shown in blue, homozygous mutant (*p404/ p404,* n=36 larvae) shown in red. F) Sensitivity index for *escapist* line sensitivity curves shown in 1E using *p404* for genotyping. Kruskal-Wallis test performed with Dunn’s test for multiple comparisons. WT vs. Mut: p<0.0001; Het vs Mut: p<0.0001; WT vs Het: p=0.8723

To identify the gene causative for the *escapist* behavioral phenotype we took an unbiased, genome-wide approach to identify single nucleotide polymorphisms (SNPs) genetically linked to the *escapist* hypersensitive phenotype. For this, we pooled a group of phenotypically mutant *escapist* larvae (pool size= 50 larvae) and a separate group of phenotypically wildtype siblings (pool size= 50 larvae) derived from the same genetic cross and performed RNA sequencing analysis(18). This analysis identified Chromosome 25 to be linked to the hypersensitivity phenotype(18). To identify potentially causal mutations on Chromosome 25, we reasoned that the mutation causing the mutant phenotype would be unique to the *escapist* line. Therefore, we searched for single base pair mutations present within the *escapist* line but absent in ten other wildtype or mutant pools on which we previously performed RNA sequencing. We prioritized single base pair mutations with read percentages close to the read percentages predicted for recessive inheritance. Since the phenotype is inherited recessively (Figure 1A-1C), the causal mutation should be found in 100% of mutant reads and approximately 33% of sibling reads. Furthermore, we prioritized single base pair mutations predicted to result in missense or nonsense mutations in protein coding regions of genes. Based on these criteria, we identified SNP *chr 25: 4531251bp T>C* within the *synaptotagmin7a (syt7a)* gene as the sole candidate satisfying all our criteria (Figure 1D). Mutant reads comprised 39% (9/23) of the *escapist* wildtype sibling pool and 93% (27/29) of the *escapist* mutant pool. Finally, the SNP was not observed in any other line we had previously sequenced (0/371 reads from ten mutant, sibling, and wildtype pools). To independently confirm that the SNP (referred to as *p404*) is linked to the *escapist* phenotype, we performed the acoustic startle sensitivity assay on subsequent generations of the *escapist* line and genotyped for *p404* after behavioral testing. When compared to larvae that are genotypically heterozygous or wild type, *p404* homozygous larvae exhibit increased acoustic startle sensitivity (Figure 1E, 1F). Together, these data strongly suggest that the SNP within *syt7a* (*p404)* is tightly linked to the *escapist* hypersensitive phenotype.

### *syt7a* mutant alleles complement the *escapist* mutant allele

*synaptotagmin 7a* is a member of the synaptotagmin gene family encoding calcium sensing proteins involved in vesicle release and replenishment(25). In addition*, synaptotagmin 7* proteins have also been shown to mediate asynchronous vesicle release(26,27). Structurally, synaptotagmin proteins are composed of a transmembrane region, a linker region, and two calcium domains (C2A and C2B). In zebrafish, exon 2 of *syt7a* encodes a transmembrane domain and the last four exons encode two calcium binding domains, C2A and C2B (Figure 2A)(28). In other vertebrates, the *synaptotagmin 7* genes encodes multiple splicing isoforms, with at least three isoforms present in mouse and in humans(29). Two alternatively spliced *syt7a* isoforms are predicted in zebrafish and expression of both has been confirmed within the zebrafish retina(28). We named the isoforms *syt7aα* and *syt7aβ*, consistent with the nomenclature of *syt7a* isoforms present in mice (Figure 2B). The difference between the two protein isoforms lies in the length of the linker region connecting the transmembrane region to the calcium binding domains, with *syt7aβ* having a longer linker domain. Alignment of the *syt7aβ* splicing isoforms reveals that the linker region affected by *p404* is highly conserved across vertebrate species (Figure 2C), with the *p404* SNP located within exon 5 of *syt7a* resulting in the conversion of a highly conserved tyrosine to a histidine (Fig 2C). The *syt7aα* isoform is predicted to be unaffected by *p404*. To test whether the *escapist* hypersensitive phenotype is caused by loss of *syt7a* gene function, we used CRISPR/Cas9 genome editing to generate three independent *syt7a* mutant alleles (Figure 2D). Sequencing of the mutant alleles revealed that *syt7a^72bpinsert^ (p431)* and *syt7a^34bpdel^ (p432)* result in a net 72bp insert and a 34bp deletion within exon 2 respectively, with both resulting in a premature stop codon and truncation of the transmembrane domain. *Syt7a^wldel^ (p433)* results in a deletion of the entire *syt7a* open reading frame and is predicted to be a protein null allele (Fig. 2D). All three mutations are predicted to affect both the *syt7aα* and *syt7aβ* isoforms. To determine whether the CRISPR/Cas9 -generated *syt7a* mutants themselves exhibit changes to their baseline threshold, we performed the sensitivity assay on *p432* larvae. Compared to their wildtype siblings, homozygous *p432* mutants do not exhibit a difference in their sensitivity curves or their sensitivity index (Figure 2E,2F)

**Figure 2.**
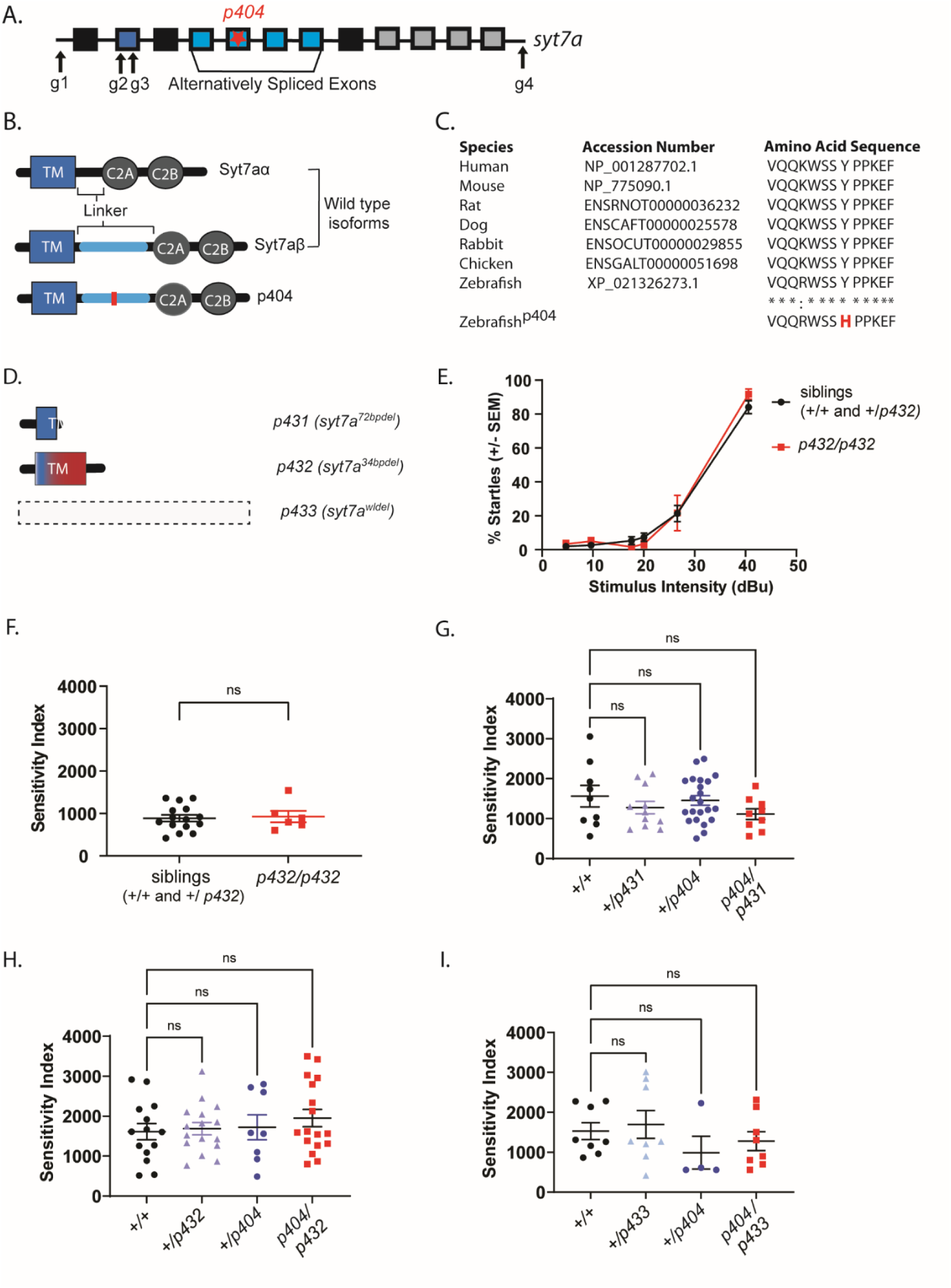
CRISPR-Cas9 generated *syt7a* mutant alleles complement the *escapist p404* mutation. A) Schematic of *syt7a* gene. Boxes indicate exons. Exon 2 (dark blue) encodes the transmembrane region of the Syt7a protein. Exons 4-7 are alternatively spliced to produce linker domains of varying lengths. The last four exons encode for the two calcium binding domains, C2A and C2B. Star represents the location of the *p404* lesion within the *syt7a* gene. Arrows indicate regions that were targeted with CRISPR guides for generation of independent mutant alleles. B) Schematic illustrating the protein structures of two *syt7a* splicing isoforms: *syt7aα* and *syt7aβ* as well as the protein structure resulting from the *p404* base pair change within the *escapist* line. The red line indicates where the missense mutation is located within the *syt7aβ* isoform. C) Clustal Omega alignment of the Syt7β protein isoform across different vertebrates. Asterisks (*) represent amino acids that are fully conserved across vertebrates, whereas colons (:) represent amino acids that are similar. Zebrafish^p404^ results in an amino acid change of a highly conserved Tyrosine (Y) to a Histidine (H) in the *syt7a*β isoform. D) Depiction of predicted protein structures from CRISPR-Cas9-generated *syt7a* mutant lines. *syt7a^72bpinsert^ (p431)* and *syt7a^34bpdel^ (p432)* encode premature stops resulting in a predicted truncated protein of both wildtype isoforms of *syt7a*. The entire *syt7a* locus is deleted in *syt7a^wldel^ (p433)*, which is predicted to produce no protein in the resulting mutant line. E) Sensitivity curve for both siblings (n=15 larvae, black line) and *p432* homozygous mutant larvae (n=6 larvae, red line). X-axis represents the stimulus intensity presented to larvae; Y-axis represents the average percent startles performed by larvae in each group. F) The area under the curve calculated from the sensitivity curve of both siblings and *p432* homozygous mutant individual larvae and plotted as the sensitivity index (AU). Mann-Whitney test was used to compare siblings to *p432* mutants (p=0.9257) G) Sensitivity Index for complementation testing between *escapist* line and *syt7a^72bpinsert^ (p431)* allele. Homozygous WT (+/+, n=9 larvae), +*/p431* (n=10 larvae), +/*p404* (n=22 larvae), *p404 / p431* (n=9 larvae). Groups were compared by using a one-way ANOVA with Tukey’s test for multiple comparisons. H) Sensitivity Index for complementation testing between *escapist* line and *syt7a^34bpdel^ (p432)* allele. Homozygous WT (+/+, n=14 larvae), +*/p432* (n=16 larvae), +/*p404* (n=8 larvae), *p404 /p432* (n=17 larvae) Groups were compared by using a one-way ANOVA with Tukey’s test for multiple comparisons I) Sensitivity Index for complementation testing between *escapist* line and *syt7a^wldel^ (p433)* allele. Homozygous WT (+/+, n=8 larvae), +*/p433* (n=8 larvae), +/ *p404* (n=4 larvae), *p404 /p433* (n=8 larvae). Groups were compared by using a Kruskal-Wallis test with Dunn’s test for multiple comparisons.

Finally, we crossed the CRISPR/Cas9 derived *syt7a* mutant alleles to the *escapist* line and assessed startle sensitivity on 5dpf larvae. In all three cases, larvae trans heterozygous for *p404* and a CRISPR-Cas9-generated allele did not exhibit a higher sensitivity index compared to their wildtype or singly heterozygous siblings (Figure 2G-I). Thus, the *syt7a* mutant alleles genetically complement the *p404* allele. This data further suggests that while the *SNP p404* in the *syt7a* gene is tightly linked to the *escapist* phenotype, loss of *syt7a* function is not causative of the *escapist* hypersensitivity phenotype.

### Zebrafish *syt7a* is dispensable for larval acoustic, visuomotor, and visual behavior

Across various species, including mice, birds, and zebrafish, *syt7* is expressed ubiquitously and is enriched in the brain. Specifically, *syt7* is expressed in auditory hair cells and the retina and moreover, *syt7* knockout mice exhibit changes in auditory pre-pulse inhibition and anxiety-like behaviors, suggesting a potential role for *syt7* in acoustically and/or visually-evoked behaviors (28,30–32). To test whether *syt7a* regulates auditory or visually evoked behaviors in zebrafish, we subjected *syt7a^34bpdel^ (p432)* mutants to previously validated behavioral paradigms including the visuomotor response(33), the dark flash response(34), the light flash response(34), and the acoustic startle response(35). In all acoustic assays, *p432* homozygous mutant larvae performed indistinguishably from their wild type siblings (Figure 2E, 2F) (Figure 3Ai, 3Aii) (Supplemental Figure 1), suggesting that both baseline and acute regulation of the acoustic startle response in *syt7a* mutants is intact. Furthermore, we detected no difference in baseline response or kinematic response to dark flashes (Figure 3Bi-3Biii), or for any tested parameters of the visuomotor response, the dark-flash response, and the light-flash response (Supplemental Figure 1). Taken together, our data indicates that in 6dpf larvae *syt7a* is dispensable for a broad range of acoustic, visual and visuomotor behaviors.

**Figure 3.**
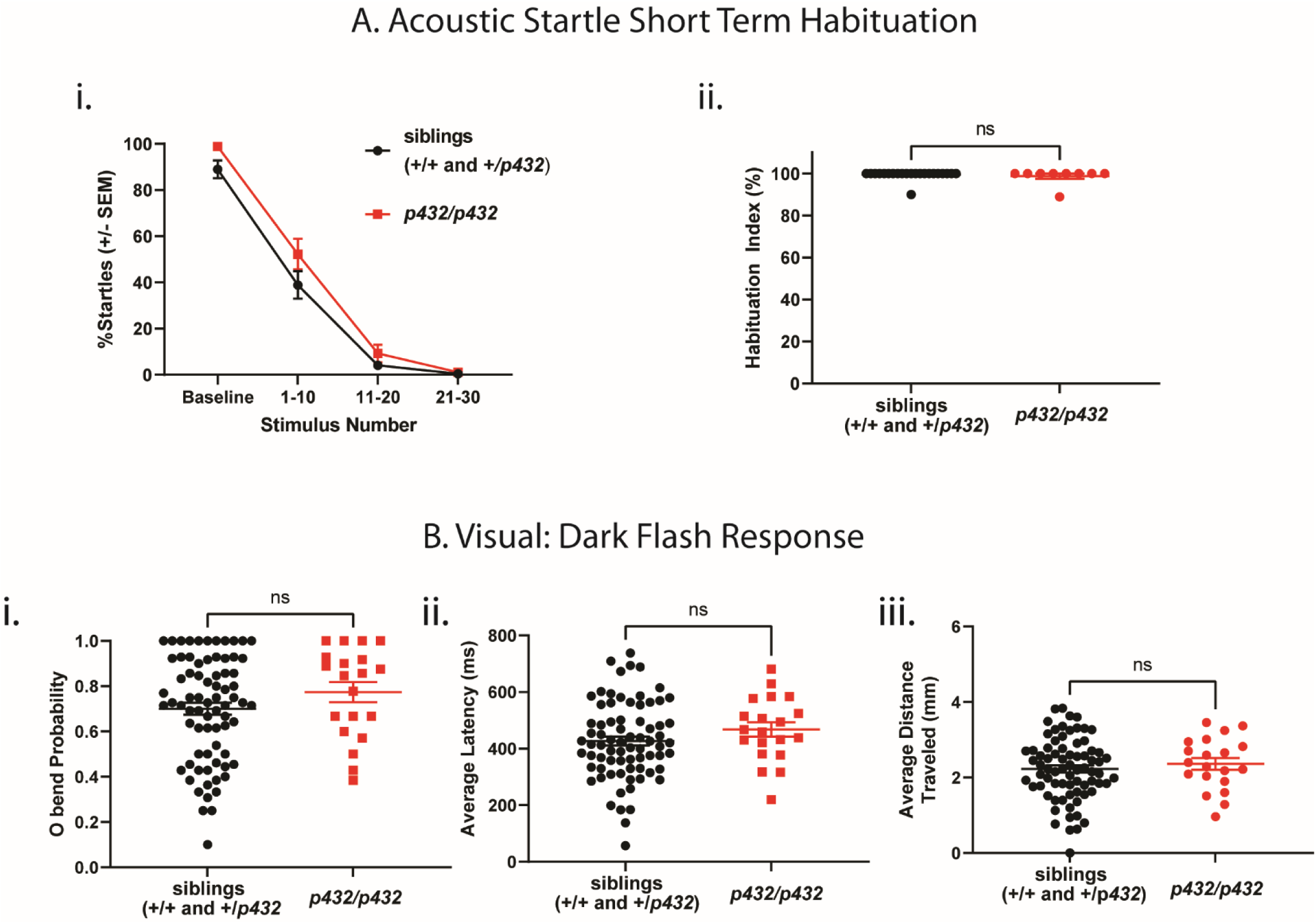
*syt7a* mutant larvae do not exhibit altered behavioral phenotype at 6 days post fertilization. A) Acoustic startle short term habituation assay. i) Percent startle of both siblings (n=22 larvae, black) and *p432* homozygous mutant larvae (n=9 larvae, red) at baseline (10 acoustic stimuli with 40 sec interstimulus interval), as well as during stimuli that induce habituation (acoustic stimulus presented with 1sec interstimulus interval). Average percent startle is reported for stimulus number 1-10, 11-20, 21-30. Any larvae that had an average probability of startle of less than 70% during baseline stimuli were omitted from habituation analysis. ii) Habituation index calculated for both siblings and *p432* homozygous mutant larvae. Habituation index is calculated using the following formula: % Habituation= [1-((%Startle Stimulus 21-30)/(%Startle Baseline))] *100. Mann-Whitney test was used to compare siblings to *p432* mutants (p=0.7871) B) Analysis of dark flash response for sibling larvae (n=74 larvae) and *p432* homozygous mutant larvae (n=20 larvae). i) O-bend probability, ii) average O-bend latency and iii) average distance traveled during O-bend are shown. For statistical purposes, values were normalized to sibling responses, and student’s t-test was performed to compare mutants and sibling groups with Bonferroni for multiple comparisons of 86 parameters analyzed during behavioral testing (See supplemental figure I for all parameters tested).

### *escapist p404* larvae exhibit baseline changes to visual stimuli

Finally, we asked whether behavioral deficits in *escapist* mutants were limited to the developmental, innate startle threshold, or whether *escapist* mutants also exhibited deficits in acute threshold regulation. Using *p404* as a genotyping marker for *escapist* mutants, we utilized an array of behavioral assays including previously validated acoustic and visual behavioral paradigms to test for the presence of additional behavioral phenotypes in *escapist* mutants(36). Consistent with our previous findings, *escapist* larvae homozygous for *p404* exhibit acoustic startle hypersensitivity as indicated by an increased sensitivity index (Figure 1E, 1F, Supplemental Figure 1), yet display no difference in acute regulation of the startle response as assayed by pre-pulse inhibition (Supplemental Figure 1) and habituation (Figure 4Ai, 4Aii). While *p404* homozygous mutant *escapist* larvae are not significantly different after Bonferroni correction in dark flash parameters such as in the probability of response to the dark flash or average latency in response (Figure 4Bi, 4Bii), we did observe significant decreases in the average distance traveled or in the average displacement during dark flash responses (Figure 4Diii, Supplemental Figure 1). Given that *escapist* mutants exhibit wildtype-like kinematic parameters of the acoustic startle response(18), this suggests differential roles for *escapist* in regulating baseline responses to acoustic compared to visual behaviors. Additionally, we do not observe differences in dark flash habituation in *p404* homozygous mutant *escapist* larvae (Supplemental Figure 1), suggesting that visual habituation remains intact. Overall, detailed behavioral characterization reveals that rather than exhibiting a global increase in responsiveness, *escapist p404* larvae exhibit an increased response probability selectively to acoustic stimuli. Importantly, changes were only observed in baseline responsiveness but not in assays that measure acute threshold regulation, underscoring the selectiveness of the behavioral phenotypes observed in *escapist p404* larvae, and hence the specificity of the process disrupted in these animals.

**Figure 4.**
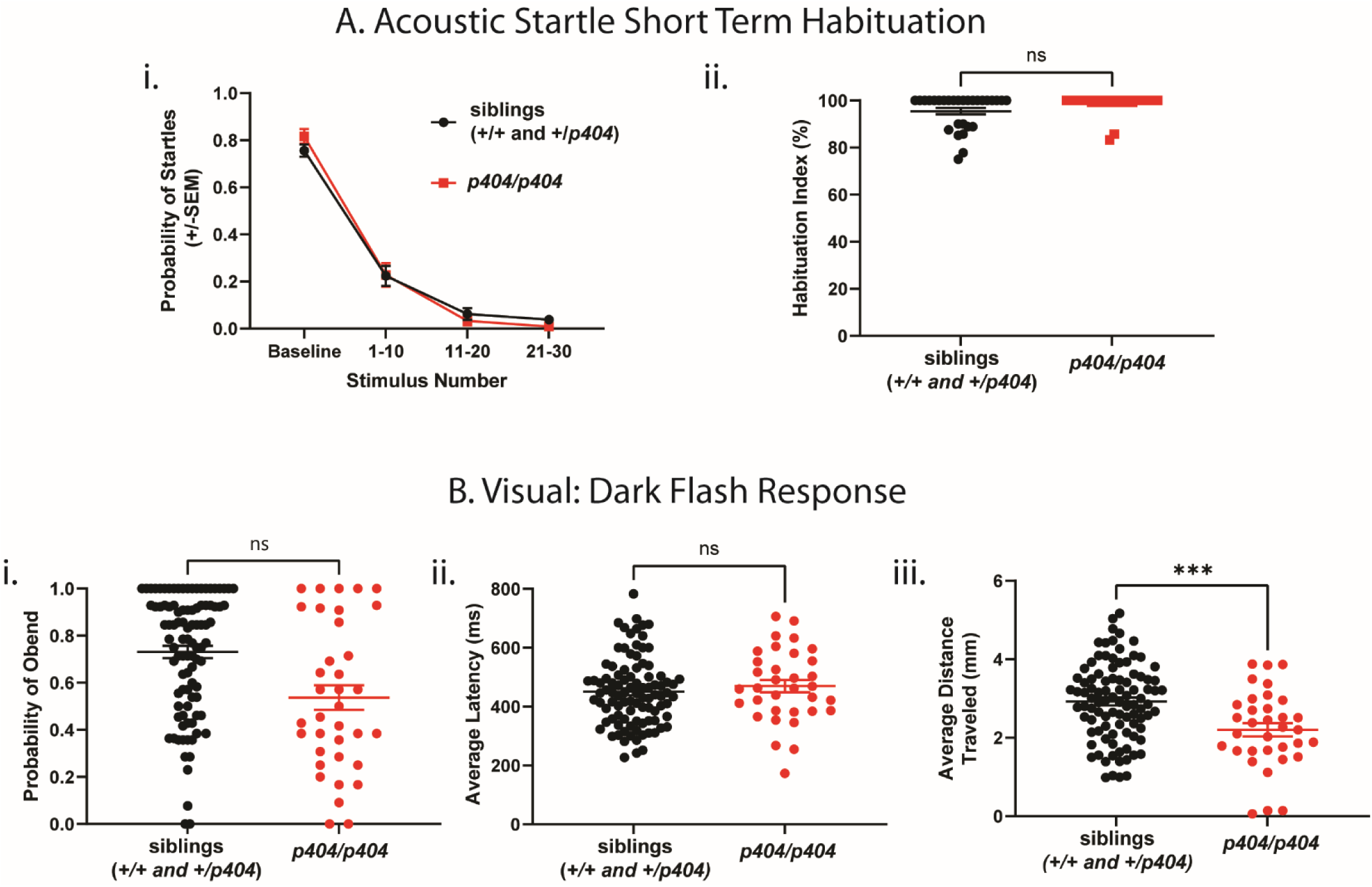
*escapist p404* larvae exhibit increased acoustic startle sensitivity and decreased visual responses at 6 days post fertilization. A) Acoustic startle short term habituation assay: i) Average probability of startle for siblings (n=29 larvae, black) and *p404* homozygous mutant larvae (n=23 larvae, red) in *escapist* line at baseline (10 acoustic stimuli with 40 sec interstimulus interval), as well as during stimuli that induce habituation (acoustic stimulus presented with 1sec interstimulus interval). Average probability of startle is grouped by stimulus number 1-10, 11-20, 21-30. Any larvae that had an average probability of startle of less than 0.7 during baseline stimuli were omitted from habituation analysis. ii) Habituation index calculated for both siblings and *p404* homozygous mutant larvae. Habituation index is calculated using the following formula: % Habituation= [1-((%Startle Stimulus 21-30)/ (%Startle Baseline))] *100 B) Dark flash response for sibling larvae (n=97 larvae) and *p404* homozygous mutant larvae (n=36 larvae). i) O-bend probability, ii) average distance traveled and iii) average distance traveled are shown. For statistical purposes, values were normalized to sibling responses, and student’s t-test was performed to compare mutants and sibling groups with Bonferroni used for multiple comparisons of 86 parameters analyzed during behavioral testing (See supplemental figure I for all parameters tested). P-values that are significant after analysis and statistical correction are indicated with asterisks (***).

## Discussion

From a forward genetic screen we had previously identified several mutant lines that display hypersensitivity to acoustic stimuli. We showed that one mutant line harbors a mutation in the cytoskeletal regulator *cyfip2*(18). Here we focus on a second mutant line from the genetic screen, the *escapist* line. Through RNA sequencing analysis, we identified a SNP (*p404*) that is tightly linked to the *escapist* mutant phenotype. *p404* is located within the *syt7a* gene, a zebrafish homolog of the *synaptotagmin 7 (syt7)* gene. Through genetic complementation, we determined that loss of *syt7a* is likely not causative of the *escapist* hypersensitive phenotype. However, by using *p404* as a tool to select for *escapist* mutants, we identify that *escapist* larvae exhibit baseline changes to auditory and visual behaviors, but not in acute regulation of these behaviors.

Synaptotagmin 7 plays roles in vesicle release and fusion(37), long term potentiation, and synaptic facilitation(38,39). *Syt7* knock out mice exhibit several behavioral deficits, including a deficit in auditory pre pulse inhibition(32). In *syt7a* mutant zebrafish larvae, we failed to detect any changes in auditory or visual behaviors, although it is possible that deficits were masked by genetic redundancy or compensation. In fact, two *syt7* paralogs, *syt7a* and *syt7b,* have been identified in zebrafish. Morpholinos targeting *syt7b* have effects on asynchronous release of vesicles in motor neurons.(40) Yet, whether *syt7b* can partially compensate for the loss of *syt7a* remains unclear. In *drosophila* loss of *syt7* has a dosage specific effect on synaptic vesicle release, consistent with the idea that expression levels of *synaptotagmin* 7 are important for its function(41). Thus, future studies involving the analysis of double mutant combinations of the synaptotagmin family members will enable further exploration of the role of *syt7* in regulating zebrafish behavior.

Through genetic complementation and behavioral testing, we determined that the *escapist* acoustic hypersensitivity phenotype is not due to loss of *syt7a*. It is possible that *p404* does not result in a loss of *syt7a* function. For example, rare, recessive antimorphic mutant alleles can result in multiple copies of the mutant allele interfering with the function of related genes and proteins(42). Knockdown of the mutant *syt7a* protein in *escapist p404* mutants could determine if the *escapist* phenotype is caused by a recessive antimorph. Alternatively, it is also possible that *syt7a* is not the causative gene. For example, SNPs in potentially causative genes that are expressed at low levels at 5-6 dpf, the developmental age at which we collected our larval samples for RNA sequencing, would not have been included within our results. Alternatively, mutations resulting in nonsense mediated decay would reduce expression of the causative gene in *escapist* mutants and preclude detection through our pipeline. Future work analyzing gene expression changes between *escapist* mutants and siblings could identify the causative gene. Furthermore, analysis of gene expression changes in *escapist* mutants could provide insight into molecular pathways affected, establishing an entry point to study the molecular mechanisms involved in establishing behavioral thresholds.

By utilizing *p404* as a genetic marker for *escapist* mutant larvae, we determined that the gene causing these behavioral changes in *escapist* is involved in a) establishing the baseline threshold for the acoustic startle response and b) involved in regulating kinematic features of baseline responses to dark flash stimuli. The gene causing these phenotypes in *escapist* may establish the acoustic startle threshold in wildtype larvae through cell types within the acoustic startle circuit. The Mauthner cells are reticulospinal neurons necessary and sufficient to activate contralateral motor neurons and initiate the acoustic startle response(43). Regulatory neurons which synapse onto the Mauthner cells, such as the feedforward excitatory spiral fiber neurons, are important for the establishment of the acoustic startle threshold(44). In *cyfip2* mutants, which also exhibit decreased acoustic startle threshold, spiral fiber activity and recruitment is increased during normally subthreshold acoustic stimuli(18). Calcium imaging on spiral fiber neurons at normally subthreshold acoustic levels in *escapist* larvae would determine if *escapist* larvae exhibit a similar increased activity in spiral fiber neurons. Alternatively, calcium imaging or activity mapping of other cell types within the startle circuit, such as auditory hair cells which provide acoustic input to the Mauthner cells, or feedforward glycinergic neurons which synapse onto the Mauthner, would allow identification of cell types that may be affected in the *escapist* line. Determining the cell types affected in the *escapist* line will help provide insight into the cellular mechanisms involved in establishing the acoustic startle threshold.

Our work has identified and characterized a useful genetic marker for identifying acoustically hypersensitive mutants within the *escapist* zebrafish line. Additionally, by utilizing this marker we have identified that *escapist* mutants show changes to establishment of acoustic and visual behaviors, while acute regulation of these behaviors in *escapist* mutants remain intact. Together, our findings support the idea that molecular and circuit mechanisms that regulate establishment of the acoustic startle response and dark flash responses can be largely independent of the mechanisms that acutely regulate these behaviors. Overall, our work lays the groundwork for determining the molecular and circuit pathways involved in establishment of baseline thresholds and furthers our understanding of the differences between acute regulation and innate behavioral thresholds.

## Methods

### Zebrafish Husbandry

#### Larvae were raised at 28-29 degrees Celsius on a 14-hour light: 10-hour dark cycle in E3 media. RNA sequencing analysis

RNA sequencing data for the *escapist* allele collected in Marsden et al, 2018 was further analyzed to identify candidate mutations for the *escapist* mutant line in the linked region Chromosome 25(18). Given that the forward genetic screen was performed using ENU mutagenesis, we focused on single base pair mutations within Chromosome 25 for potential candidate mutations. Causative mutations are expected to be recessive and therefore are expected to comprise close to 100% of reads within the *escapist* mutant pool and approximately 33% in the sibling pool. Priority was given to mutations that were predicted to have deleterious changes to coding sequences of genes, such as nonsense, missense mutations, or changes to splicing donor or acceptor sites.

### Generation of CRISPR derived *syt7a* alleles

Guides targeting *syt7a* were designed by using CHOP CHOP and IDT software. Guides were designed to target exon 2, which encodes the transmembrane domain of *syt7a* protein (guides 2 and 3), as well as the 5’ and 3’ UTR (guides 1 and 4). WIK wildtype embryos were co-injected with either guides 2 and 3, or guides 1 and 4 for a whole locus deletion. Injected G0 embryos were raised to adulthood. G0 fish were then outcrossed, F1 larvae were collected and screened to identify mutant carriers. F1 mutant larvae were raised to adulthood, outcrossed and F2 larvae grown to generate stable mutant lines. F2 carriers from each stable line were then incrossed and gDNA was collected from F3 homozygous mutants. F3 gDNA was then amplified using primers flanking the CRISPR targeted regions, TOPO cloned, and Sanger sequenced to confirm each mutation.

### Behavioral analysis

Zebrafish larvae were behaviorally tested between 4-6 dpf.

### Acoustic Startle Hypersensitivity Assay

Larvae were presented with 6 different stimulus intensities, each presented 5-10 times with an interstimulus interval of 40 seconds. Larval responses to the stimuli were recorded using a high-speed camera (Photron Fastcam MiniUX) at 1000 frames per second. Videos were then tracked using FLOTE v2.0 and Batchan to analyze behavior of each individual larvae as previously described(24). The average percent of escape responses performed by larvae was recorded at each stimulus intensity. The area under each curve was then calculated using PRISM software to obtain the sensitivity index.

### Acoustic Startle Habituation Assay

Larvae were presented with 40 acoustic stimuli total. During the pre-habituation phase, larvae were presented with 10 stimuli with a 40 second interstimulus interval. During the habituation phase, larvae were presented with 30 stimuli with a 1 second interstimulus interval. Stimuli were grouped into 4 different bins: baseline response, habituation stimuli 1-10, habituation stimuli 11-20, habituation stimuli 21-30. The average percent of larvae responding with escape responses were recorded during each habituation bin. Larvae that did not respond with over 80% startles at baseline were omitted from further analysis. Habituation index is calculated using the following formula: % Habituation= [1-(%Startle Stimulus 21-30)/(%Startle Baseline)] *100

### Complementation testing

Independently generated *syt7a* heterozygous F2 fish were crossed to *escapist* carriers heterozygous for the *p404* single base pair mutation. Larvae were raised to 5dpf and tested on the acoustic hypersensitive assay. Larvae were collected after testing and genotyped for the independently generated *syt7a* mutant allele and for *p404*.

### Broad behavioral assay at larval stages

Broad behavioral assay at 6dpf larval stage was performed as described in Campbell et al, 2023(36). Briefly, 6 days post fertilization larvae were placed in individual wells in a 10×10 well acrylic testing arena. The following behavioral tests were performed in the following order: Visual Motor Response (Light ON followed by Light OFF), Light Flash, Dark Flash, Acoustic Startle Response Behaviors. Approximately 50 larvae from each cross were tested per trial. Larvae were genotyped after behavior was performed. A total of 86 parameters were tested across the various behavioral paradigms. For statistical purposes, values were normalized to sibling responses, and student’s t-test was performed to compare mutants and sibling groups with Bonferroni used for multiple comparisons of 86 parameters analyzed during behavioral testing.

#### Visuomotor response

Larvae were placed in the testing arena and allowed to acclimate for 30 minutes before testing. Larvae were then recorded for 8 minutes in light (Light ON), and 8 minutes in the dark (Light OFF). Video was recorded at 20 frames per second.

#### Light Flash

Larvae were presented with 15 500ms white light stimuli after previously being in dark conditions. Each stimulus was separated by an interstimulus interval of 30 sec. Responses were recorded at 500 frames per second.

#### Dark Flash Baseline

Larvae were presented with 15 1sec dark flash stimuli after previously being in light conditions for 5 minutes. Each stimulus was separated by 30 sec interstimulus intervals. Responses were recorded at 500 frames per second.

#### Dark Flash Habituation

Larvae were presented with 4 blocks of 14 1 sec dark flash stimuli after Dark Flash baseline stimuli. Dark flash habituation stimuli were separated by 10 second interstimulus intervals. The first and third block of dark flash habituation stimuli were unrecorded, while the second and fourth block of dark flash habituation were recorded at 500 frames per second.

#### Acoustic Behaviors

Larvae were presented with the following stimuli (recorded at 500 frames per second): Pre-pulse Inhibition: low intensity pulse followed by high intensity pulse with inter-stimulus interval of 50ms, High intensity stimuli (10 times), Acoustic habituation: High intensity acoustic stimuli delivered 30 times with 1 sec ISI.

#### Statistical Analysis

Graph generation and statistical analysis was performed using a combination of Python as well as Graphpad Prism (www.graphpad.com). The D’Agostino and Pearson test was used to assess normality. If data was not normally distributed, the Mann-Whitney or Kruskall-Wallis test with Dunn’s test for multiple corrections was used to compare groups. If data was normally distributed, the students t test or one-way ANOVA with Tukey’s test for multiple corrections was used to compare groups.

## Acknowledgements

The authors would like to thank the Granato lab members for their feedback on our findings and on the manuscript. The authors would also like to acknowledge the University of Pennsylvania Genomic Analysis Core DNA sequencing Facility. This work was supported by grants to M.G (NINDS R01NS118921). EAO was supported by NIDCD 5T32DC016903. P.D.C. was supported by T32MH019112 and R25MH119043. J.C.N. was supported by NINDS K99NS111736 and NINDS R00NS111736.

## Author Contributions

Conceptualization: E.A.O. and M.G.; Investigation: E.A.O., J.C.N., P.D.C.; Data Curation: E.A.O. and P.D.C.; Formal Analysis: E.A.O., J.C.N. and P.D.C.; Funding Acquisition: E.A.O. and M.G.; Resources and Supervision: M.G.; Writing – Original Draft: E.A.O.; Writing-Review and Editing: E.A.O, P.D.C., J.C.N., M.G.

