## Supplemental Figure 1 for "A single base pair substitution on Chromosome 25 in zebrafish distinguishes between development and acute regulation of behavioral thresholds"

**Supplemental Figure 1. Comprehensive behavioral analysis of escapist *(p404)* and *syt7a*$^{34bpdel}$ *(p432)* at 6 days post fertilization**

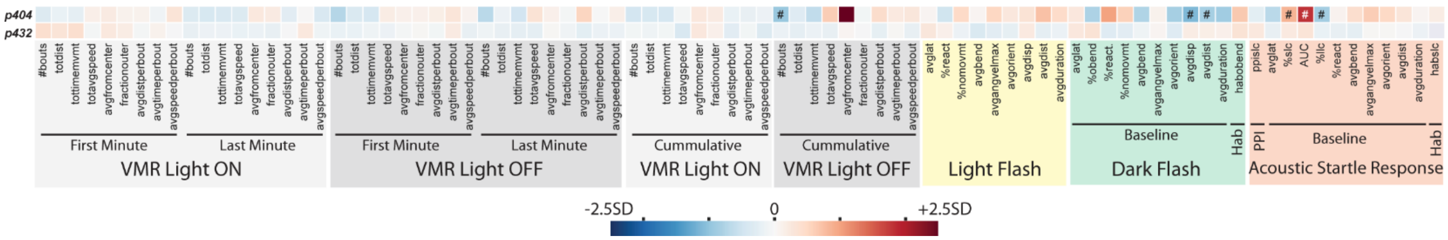


a) Heat map summary of the results from our broad behavioral assay for *escapist (p404)* line and *syt7a*$^{34bpdel}$ *(p432)* line. Different behavioral assays are designated by block: VMR Light ON (light gray box), VMR Light OFF (dark gray block), Light Flash (yellow block), Dark Flash (green block), and Acoustic Startle Response (red block). Each box within a block represents a different parameter tested during each assay. Responses from homozygous mutants were normalized to responses from their respective siblings (WT and Hets combined). The colors of the heatmap represent the Z-score, or the difference between the normalized average response from homozygous mutants compared to the average responses of their respective siblings in standard deviations, with red signifying mutants had a larger response than siblings and blue signifying mutants had a smaller response than siblings. A student t-test was used with a Bonferroni correction to correct for the multiple comparisons performed in the broad behavioral assay. Parameters that had a significant p-value after the Bonferroni correction are indicated with the pound sign (#) within the box.
